# Directional motor encoding captured in human micro-ECoG enables brain–computer interface cursor control

**DOI:** 10.64898/2026.11.13.766241

**Authors:** Kurt R. Lehner, Rebecca J. Greene, Shiyu Luo, Daniel Candrea, Ramin Anushiravani, Yoon Woo Byun, Joshua S. Miller, Mark Murphy, Jacqueline Dister, Katrina Barth, Aneil Srivastava, Elton Ho, Esha Ghorpade, Jack Berkowitz, Federico Claudi, Molly Lucas, Craig H. Mermel, Jayme Strauss, Saadi Ghatan, Christopher P. Kellner, Joshua B. Bederson, Khalil Husari, Jordina Rincon-Torroella, Youssef Comair, Chetan Bettegowda, Benjamin I. Rapoport, Nathan E. Crone

**Author notes:** These authors contributed equally to this work (Kurt R. Lehner, Rebecca J. Greene).

## Abstract

Brain–computer interfaces translate neural activity into control signals, restoring autonomy for people with paralysis. Electrocorticography, in which thin electrodes rest on the cortical surface without penetrating the brain, offers a promising path to long-term neural interfaces, yet prior surface recordings have not achieved continuous, intuitive directional cursor control. Prior approaches required indirect decoding strategies and achieved only modest performance, leading to a prevailing view that surface field potentials encode only movement speed, but not direction. Here we show that a high-density thin-film micro-electrocorticography array captures direction-specific population dynamics in human hand motor cortex that support real-time closed-loop two-dimensional cursor control after minutes of training. Moreover, dimensionality reduction and permutation analyses show that directional encoding depends jointly on the temporal evolution and spatial coordination of distributed population activity. These findings overturn the view that surface recordings lack directional content and extend dynamical-systems models of motor control, previously restricted to intracortical recordings, to mesoscopic population responses at the cortical surface, establishing a platform for the study of human motor computation and the development of clinical neural interfaces.

## 1 Introduction

Voluntary movement is governed by the coordinated dynamics of neural populations in motor cortex, yet restoring this control for people who have lost it remains a fundamental challenge[1–6]. Brain-computer interfaces aim to bridge this gap by translating cortical activity into control signals. Electrocorticography (ECoG), a neural recording technique in which surface electrodes rest on the brain without penetrating it, offers an attractive path to long-term implants as they avoid the surgical risks and inflammatory tissue responses associated with penetrating devices [7–17]. However, prior ECoG studies have not achieved the continuous, intuitive directional control that practical daily use demands. Existing approaches have relied on indirect strategies, such as mapping discrete imagined actions to independent cursor axes, and have achieved only modest performance [5, 6, 18–25]. This has led to a prevailing view that cortical surface field potentials encode movement speed rather than direction, with robust directional tuning believed to require intracortical neural recordings [26, 27]. Whether the cortical surface, sampled at sufficient spatial density, contains the directional information needed for high-performance control has remained unresolved.

Here, we show that a high-density thin-film micro-ECoG array placed over the human hand motor cortex resolves this limitation. Recording from five participants during intraoperative and short-term postoperative sessions, we demonstrate that neural population activity recorded from the cortical surface supports real-time closed-loop two-dimensional cursor control after minutes of decoder training, with performance sufficient for practical use. Dense sampling of a compact cortical region reveals that directional information is jointly encoded in the temporal evolution and spatial coordination of distributed population activity rather than a directional somatotopy, extending dynamical-systems models of motor control to the cortical surface [28–31]. These findings establish high-density surface recording as a scalable platform for both the study of human motor cortical computation and the development of clinical brain–computer interfaces, and introduce an intraoperative framework for the rapid evaluation of neural decoders in humans.

## 2 Results

To test whether high-density surface recordings could support continuous directional cursor control, we recorded local field potentials from a micro-ECoG array placed over hand motor cortex in five participants during two-dimensional target-acquisition tasks (Fig. 1). Participants first controlled cursor velocity with a contralateral joystick, then performed blocks in which cursor motion was driven by real-time decoding of neural activity. Four participants (P1–P4) were recorded intraoperatively during awake craniotomy for tumor resection, with each session limited to a brief period of joystick-controlled training followed by neurally controlled cursor use. This intraoperative cohort also served as a development cohort in which decoding approaches were systematically refined. P1 and P2 used a directional classifier, while P3 and P4 used continuous velocity decoding informed by analysis of data from earlier participants. A fifth participant (P5) received short-term micro-ECoG implantation during a postoperative monitoring period, enabling recording sessions over two consecutive days and evaluation of cross-session decoder generalization and cursor control without overt movement.

**Fig. 1.**
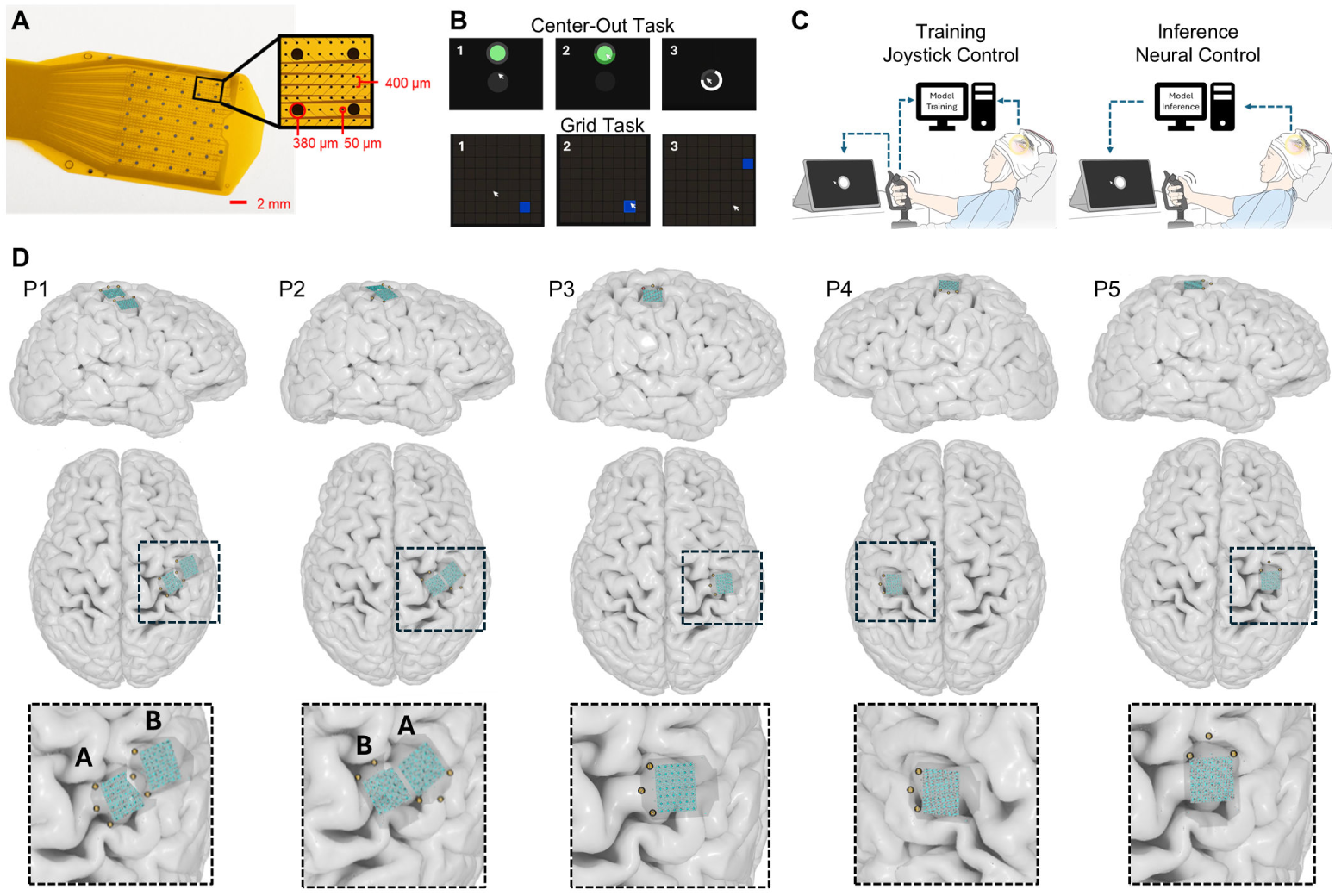
High resolution micro-ECoG and Task Design. a, Thin-film high-density micro-ECoG array used in this study, comprising 1,024 electrode contacts spanning 1.5 cm^2^, with a 400 *µ*m pitch and electrode diameters ranging from 50–380 *µ*m, fabricated on a flexible polyimide substrate. b, Behavioral tasks used for neural decoding. In the center-out task, P1-3 moved a cursor to a series of subsequent circular targets, each spaced by a fixed length vector oriented towards one of four or eight principal directions. In the grid task, P4 and P5 moved the cursor to visually cued locations within an N×N grid (4×4 for P4, 12×12 or 30×30 for P5 training blocks, and 8×8 for most P5 inference blocks except for a few outliers at 12×12 and 4×4). c, Experimental structure. During training blocks neural activity was recorded while participants controlled the cursor using a joystick. During inference blocks cursor motion was driven in real time by micro-ECoG signals using trained decoders. In most inference blocks participants continued to modulate the joystick; in selected blocks, participants were instructed to avoid overt hand movement for the assessment of control during attempted or imagined movement (P5). d, Electrode array placement over the hand area of motor cortex for all participants (P1–P5), shown on atlas reconstructions of the cortical surface. Insets show magnified views of array placement. In participants P1 and P2, two arrays were implanted, but only array A was used for the online decoding. Offline analysis for P1 used array B, which was found to have better decoding.

### 2.1 Real-time micro-ECoG supports closed-loop cursor control

Real-time closed-loop cursor control was achieved using continuous velocity decoders trained on minutes of joystick-based task data. P3-5 used a joystick to control cursor velocity in an eight-directional center-out task (P3), or a grid target task (P4, P5). Using decoders trained on 7.73 - 40 minutes of data P3-5 performed closed-loop continuous cursor control. We evaluated closed-loop cursor control by measuring target acquisition rate and median completion time relative to joystick-based physical control during open-loop training sessions. This ratio provides a clinically interpretable, device-agnostic measure of BCI usability that jointly reflects decoding accuracy and control.

P3-P4 completed the task during awake intraoperative mapping, with decoders trained on 13.6 and 12.1 minutes of data, respectively. For P3 performance biased toward rightward movements, with 47.1% of trials completed (8 of 17). Successful target acquisition required a median 10.71s, in comparison to 6.30s for tasks completed with the joystick. P4 completed 90.6% of trials (77 of 85) with a median completion time of 3.74s (Fig. 2).

**Fig. 2.**
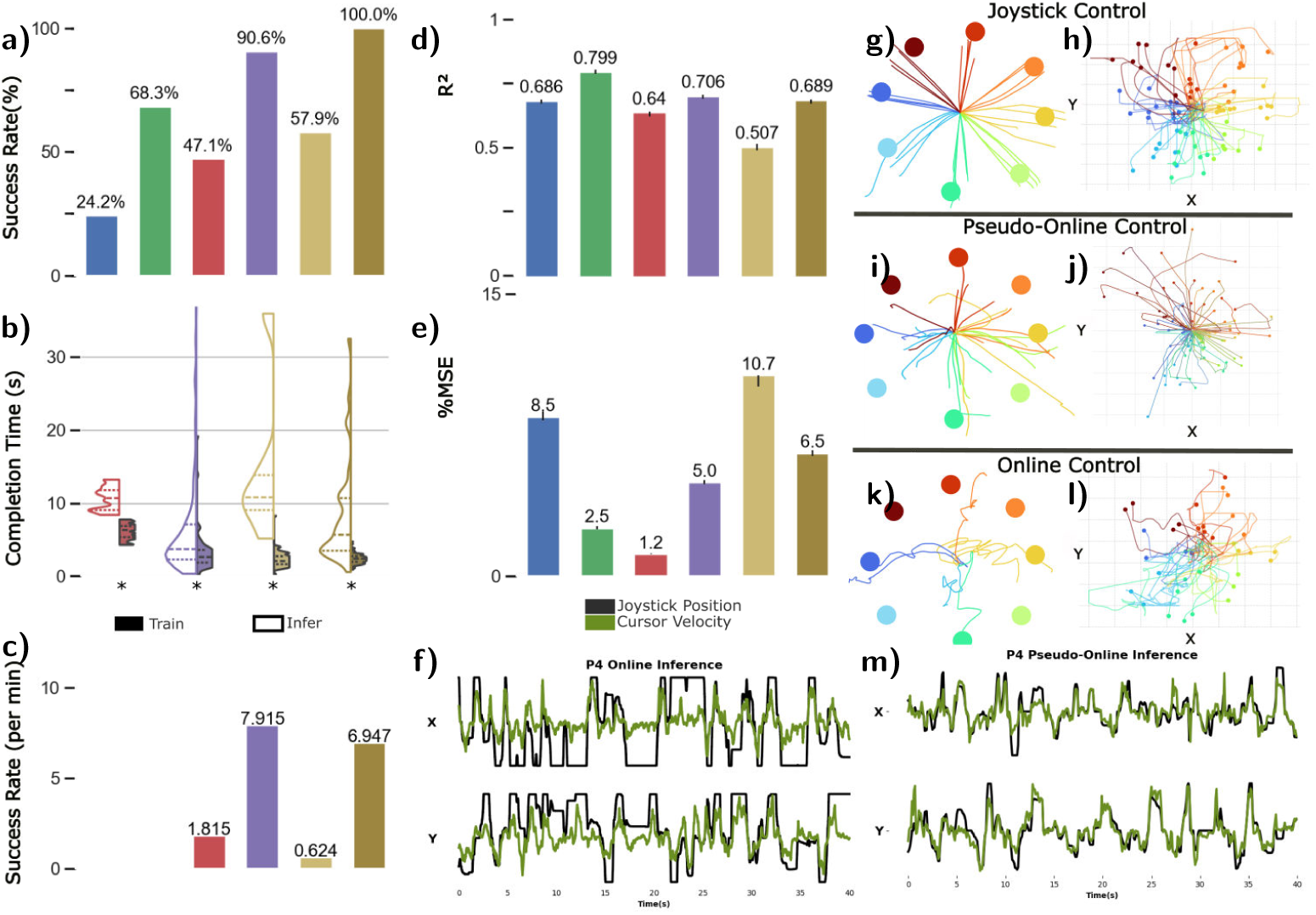
Real-time closed-loop cursor control and optimized velocity decoding from high-density micro-ECoG. Online performance metrics for each participant (P1–P5), showing a, task completion rate, b, elapsed time for successful trials during the initial inference block, and c, task throughput (see Fig 3 for P5 cross-session performance). In the intraoperative setting, P4 acquired 90.6% of targets with a median completion time of 3.74[2.30-7.07]s (joystick: 2.66[1.87-3.60]s). P3 acquired 47.1% of targets with a median completion time of 10.71[9.10-11.80] s (joystick: 6.30[5.35-6.86] s). In the extraoperative setting, P5 acquired 57.9% of targets in the initial session (10.61[8.40-24.24] s; joystick: 2.13[1.67-2.79] s) and 100% of targets one day later with a decoder trained on approximately 17 minutes of accumulated data (median 5.75[3.54-10.67] s; joystick: 2.39[2.02-2.75] s). All values are median[IQR]. Asterisks denote significant pairwise differences between blocks (two-sided Mann Whitney U test; *P ≤ 0.05, exact P values are given in Supplementary Table ??. Decoding performance for the pseudo-online analysis, showing d, coefficient of determination (*R*^2^) and e, mean squared error (MSE) between predicted and true joystick velocities across both dimensions (See Table ??). f,m Example time series of joystick-derived (black) and decoder-predicted (green) velocities during online inference and simulated inference for P4. g,h Cursor trajectories derived from joystick position during training blocks for P3 (center-out task) and P4 (grid task). Each trace represents a single trial, colored by intended movement direction. Center-out and grid trajectories have been aligned to a common starting position. i,j, Simulated control trajectories generated using participant-optimized decoders evaluated in pseudo-online fashion on held-out data. k,l, Online neural control trajectories from completed target acquisition trials.

P5 completed the tasks over a 3 day postoperative period, across which multiple model instances were trained and utilized for closed-loop cursor control (Fig. 3). In the initial session of the extraoperative study, P5 completed 57.8% of trials (11 of 18) at 10.61s after 7.73 minutes of training data collection. Less than one day later, with a decoder trained on approximately 18 minutes of accumulated data, P5 completed all 30 trials with a median time of 8.35s.

**Fig. 3.**
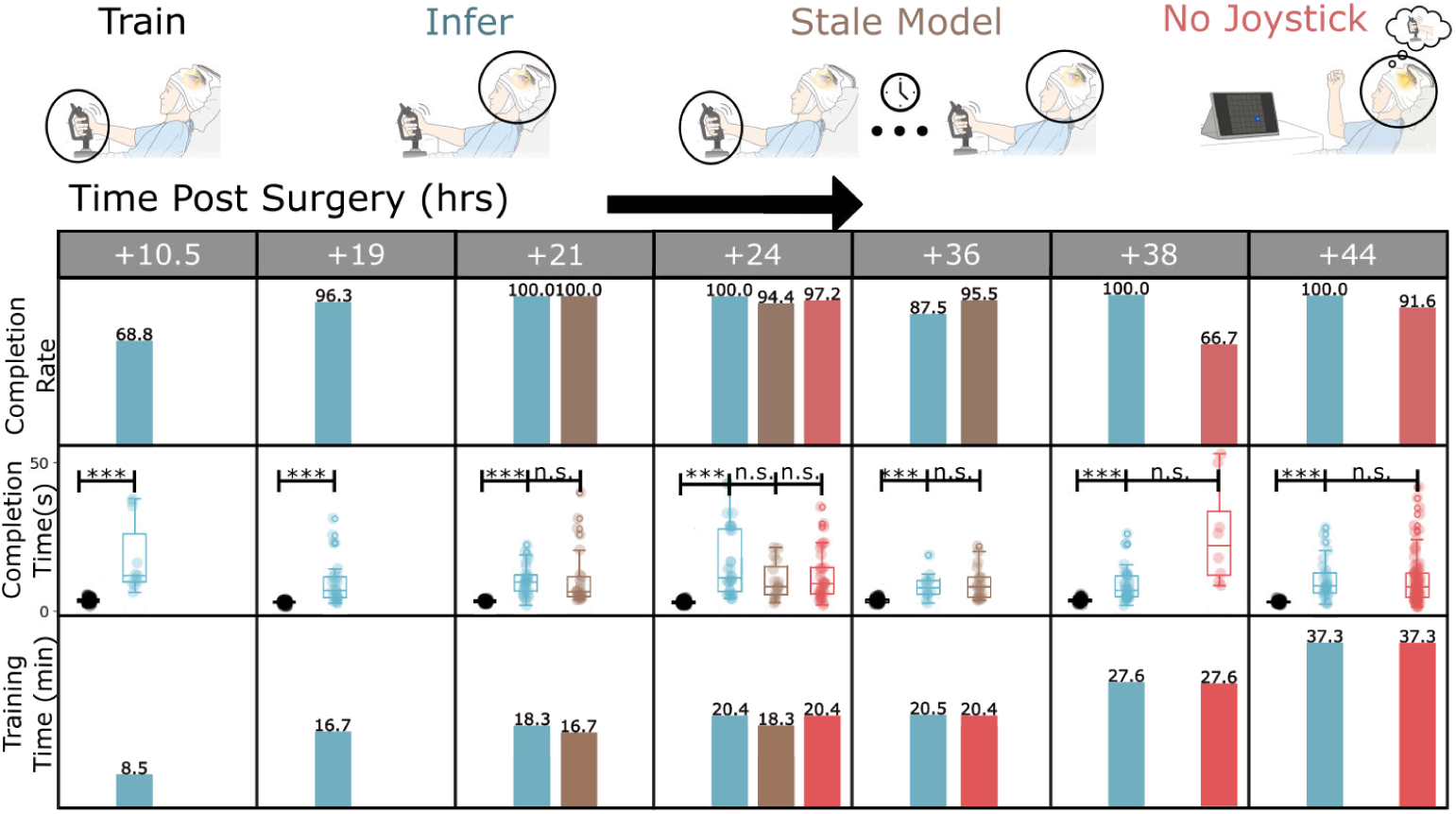
Decoder performance improves across sessions, generalizes without retraining and persists without overt movement (P5). Grid-based target task success rate and task completion time distributions are shown across postoperative recording sessions (10–44 hours). All completion times are reported as median ± s.d., with joystick-control times in parentheses. Blue markers indicate performance using a model trained during the same session: initial session, 69% target acquisition (median 10.61 ± 11.57 s; joystick: 2.13 ± 1.05 s); 24-hour session, 100% target acquisition (median 9.34 ± 5.36 s; joystick: 2.16 ± 0.72 s). Brown markers indicate performance using a model trained on the day prior without retraining: 95.5% target acquisition (median 7.24 ± 7.46 s; joystick: 2.67 ± 0.72 s). Red markers indicate performance during trials in which the participant was instructed to avoid overt hand movement: 97.2% target acquisition, 35 of 36 trials (median 8.16 ± 8.73 s; joystick: 2.23 ± 0.59 s). Accuracy improved across sessions, reaching 100% target acquisition at 24 and 44 hours postoperatively.

An initial discrete classification approach in early participants (P1-P2) yielded variable performance. A four-class directional classifier was trained on 12.58 (P1) and 12.42 (P2) minutes of task data. P1 achieved 24.2% class-weighted accuracy (9 of 37) relative to chance level of 25%, with successes largely restricted to the upwards direction. It is worth noting that although two arrays were placed for P1, only one was utilized for the online experiment. Offline analysis suggests that evaluation with the other array, positioned more laterally along the motor strip, would have improved performance. P2 achieved 68.3% class-weighted accuracy (21 of 35) relative to chance level of 25% with reliable decoding of down- and leftward movements. These results, combined with the strong directional content revealed by pseudo-online analyses, motivated the transition to continuous velocity decoding.

### 2.2 Optimized decoding reveals additional directional information in micro-ECoG signals

The closed-loop results above were achieved with decoders deployed under intraoperative time constraints, using manually selected hyperparameters without systematic validation. To assess the full directional information recoverable from micro-ECoG signals, we applied a standardized pseudo-online evaluation in which participant-specific hyperparameters were optimized on held-out data and decoders were tested causally on the validation set (See Methods). This convention was necessitated by the limited recording time and availability of labeled data. Predicted cursor velocities closely tracked joystick-derived kinematics, capturing movement onset, directional changes, and speed (Fig. 2). Across all five participants, *R*^2^ values were 0.69, 0.8, 0.64, 0.71 and 0.51, with corresponding percent mean squared error (% MSEs) of 8.5%, 2.5%, 1.20%, 5.02%, and 10.7%. These results indicate that the cortical surface signals recorded here contain substantially more directional information than was extracted during real-time control, suggesting that further decoder optimization could narrow the gap between neural and joystick-level performance.

### 2.3 Decoder performance generalizes across sessions and persists without overt movement

The extraoperative case study in P5 enabled evaluation of cross session decoder stability and control in the absence of overt movement, two properties critical for chronic BCI deployment. To assess cross-session generalization, P5 performed closed-loop cursor control using a decoder trained exclusively on data collected the prior day, with no retraining or recalibration. Despite the 12-hour interval, P5 completed 95.5% of targets on an 8×8 grid with a median completion time of 5.19s, suggesting the relationship between micro-ECoG population dynamics and motor intention was sufficiently stable across days to support reuse of trained decoders (Fig. 3). To evaluate whether control could be maintained without overt movement, a necessary feature for deployment in participants with paralysis, P5 performed neurally controlled trials without simultaneous joystick motion. P5 completed 97.2% of targets (35 of 36), providing preliminary evidence that the directional signals captured by micro-ECoG are not solely dependent on movement execution or proprioceptive feedback. This result is limited to a single participant performing a small number of trials, and confirmation using attempted or imagined movement paradigms in people with paralysis will be essential.

### 2.4 Submillimeter electrode density is required for directional decoding

A central question for clinical translation is whether the full electrode density of the micro-ECoG array is necessary, or whether sparser configurations could provide comparable performance. To address this, we simulated lower-density configurations by subsampling electrodes at systematically increasing spacings and evaluated each configuration using the pseudo-online decoding framework with spatial convolutions restricted to single electrodes to prevent the model from exploiting spatial relationships absent in sparser arrays (see Methods).

Across all five participants, performance declined with increasing pitch (Fig. 4), providing a quantitative explanation for why prior clinical ECoG systems, which sample at spacings of several millimeters or more, have not supported continuous directional control. For participants who performed center-out tasks (P1-P3), this relationship was roughly linear (*R*^2^ = 0.79, 0.84, 0.51 respectively), with a negligible difference in fit quality compared to nonlinear models, demonstrated by Akaike Information criterion values of 0.19, 2.00, and 2.00 respectively. Participants who performed the grid task (P4–P5) showed a pronounced exponential decay (R² = 0.80, 0.91), with the trend becoming notably more non-linear as task complexity (grid size) increased.

**Fig. 4.**
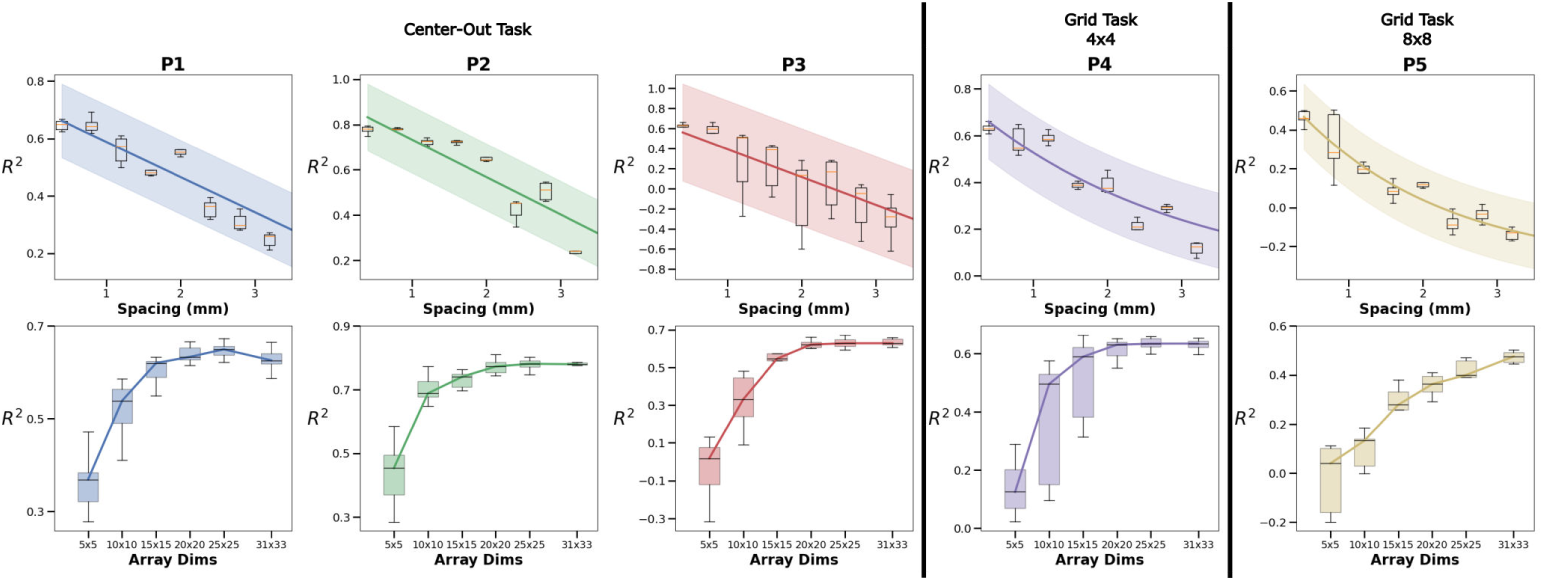
Submillimeter electrode density is required for directional decoding. Pseudo-online velocity performance (*R*^2^) is shown as a function of simulated inter-electrode spacing for participants P1-P5. a, Lower-density configurations were generated by systematically subsampling electrodes from the ≈ 1.5*cm*^2^ recording region. Each point represents mean performance across three independently trained models per configuration and shaded regions represent 95% confidence intervals. b, Electrode configurations with reduced area were generated via regular sampling of square NxN subregions within the original coverage area.

Reducing the array’s spatial footprint while maintaining full electrode density preserved decoding performance down to approximately 40% of the original coverage area for P1–P4. However, because spatially informative sites were distributed in unpredictable cortical clusters rather than concentrated in a single focal region (see below), a smaller array would risk missing critical zones entirely. The redundancy afforded by the current footprint instead confers robustness to partial channel loss, a property with direct relevance to chronic devices in which individual electrodes may degrade over time. Notably, this trend did not hold for P5, who performed tasks on grids as large as 30×30, demanding directional control across hundreds of target locations. This suggests that increased array coverage may be necessary as task complexity approaches real-world demands.

### 2.5 Directional decoding is robust to channel subsampling but depends on distributed spatial coverage

A separate question from electrode density is whether the full channel count of the array is necessary once submillimeter spacing is achieved. We retrained decoders on randomly subsampled channel sets while preserving the spatial convolution architecture (see Methods). Performance remained stable with as few as 40% of channels retained, indicating substantial redundancy in the recorded population. Removing the spatial convolution and retraining with single-electrode inputs at matched channel counts reduced performance to levels consistent with the density analysis, confirming that local spatial structure among neighboring electrodes contributes to directional decoding beyond what individual channels provide independently.

To determine whether informative channels were spatially concentrated or distributed, we performed a multi-scale spatial ablation in which contiguous electrode tiles of increasing size were sequentially zeroed during validation. Across all five participants, spatially coherent regions of high importance emerged, and the structure of these importance maps was stable across scales, indicating that directional information is organized in distributed mesoscale cortical patches rather than isolated sites. Together with the density analysis, these results establish that directional decoding depends on distributed population signals sampled at high spatial density, and that the redundancy within the current array footprint confers robustness to partial channel loss.

### 2.6 Movement direction is encoded in low-dimensional micro-ECoG population dynamics

Trial-averaged high-gamma activity revealed direction-dependent spatial activation patterns distributed across the array, but these static maps did not fully separate movement directions (Fig. 5). We therefore examined the temporal evolution of population activity using dimensionality reduction (see Methods). In low-dimensional state space, neural activity formed smooth, continuous trajectories that diverged by movement direction shortly after movement onset. A linear classifier applied to the population state at each time point confirmed that directional information increased rapidly after movement onset and peaked during movement execution (Fig. 5).

**Fig. 5.**
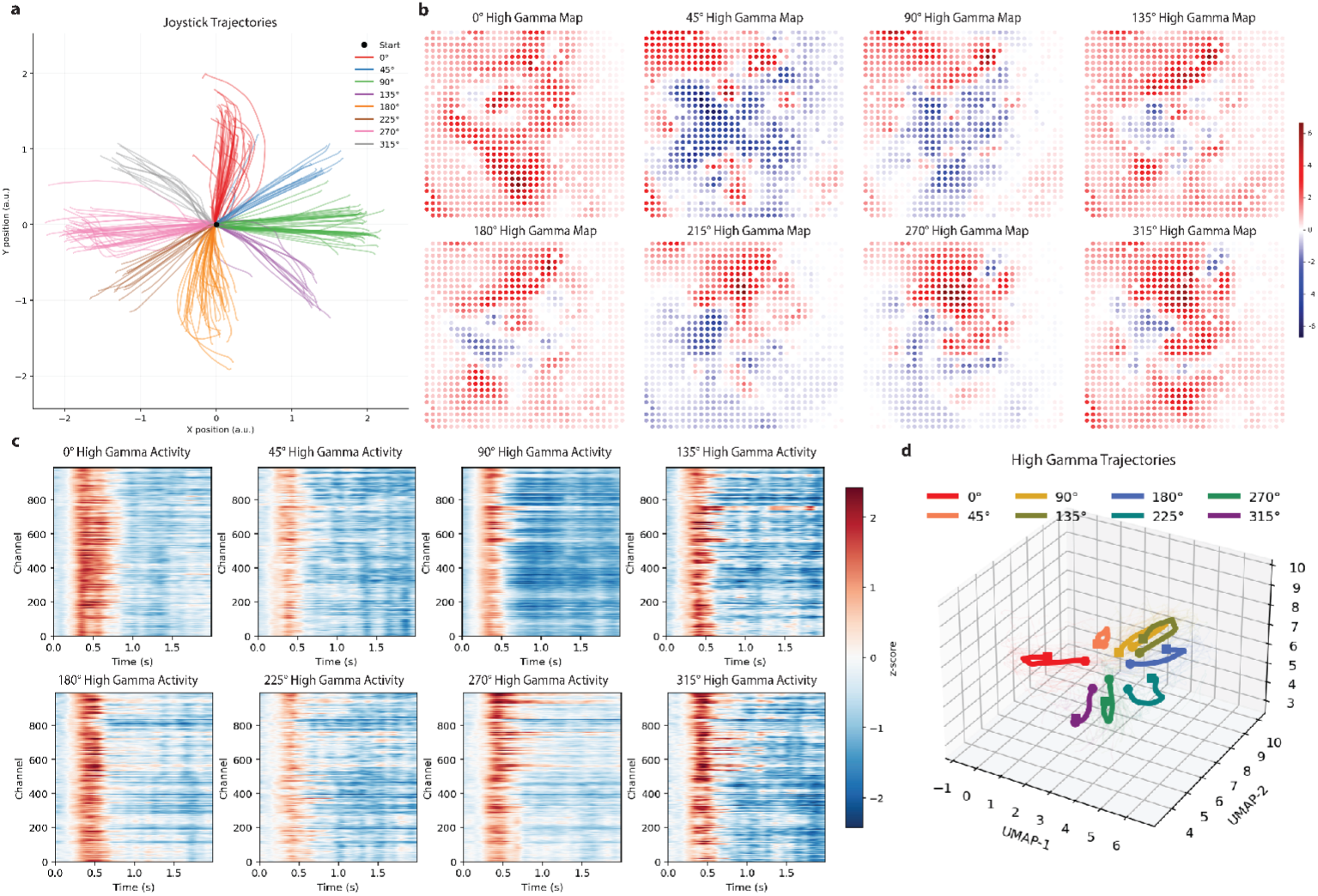
Direction-specific motor information is reflected in distributed high-gamma activity and low-dimensional population dynamics. a, Example joystick trajectories from P2 during center-out reaching tasks, colored by movement direction (0°-315°). Each trace represents a single trial aligned to movement onset. b, Corresponding trial-averaged HGA across the 1,024-channel micro-ECoG array for each reach direction, displayed as spatial maps of normalized power. Direction-dependent activation patterns are distributed across the cortical surface. c, Channel-by-time heat maps of HGA aligned to movement onset for the same directions, illustrating movement-locked temporal dynamics across the array. Rows represent channels and columns represent time. d, Low-dimensional embedding of population activity (PCA-to-UMAP). Mean neural trajectories for each direction form distinct paths in latent space, indicating separable direction-specific population dynamics.

To determine whether this directional structure depended on temporal dynamics, spatial coordination, or both, we constructed two permutation controls. Shuffling the temporal order of samples within each trial while preserving spatial correlations substantially reduced both decoding accuracy and trajectory separation. A complementary control that preserved temporal dynamics but disrupted spatial relationships across channels produced a similar degradation. Neither permutation collapsed performance entirely, indicating that each dimension retains limited directional information in isolation — but neither alone recovered the full encoding observed in the intact data. The relative contributions of temporal and spatial structure varied across participants, and overall decoding accuracy ranged from approximately 0.4 to 0.7, likely reflecting differences in electrode placement and signal quality.

Together, these analyses demonstrate that directional information is jointly encoded in the temporal evolution and spatial coordination of micro-ECoG population activity, rather than being recoverable from either dimension alone.

## 3 Discussion

We demonstrate that high-density micro-ECoG captures direction-specific population dynamics from the human cortical surface that are sufficient for real-time, continuous cursor control, a capability that prior surface recordings have not achieved with a similar-sized implant. Critically, our analyses reveal that this directional information resides in the joint temporal evolution and spatial coordination of distributed population activity. Low-dimensional embeddings of micro-electrocorticography signals formed smooth, direction-specific neural trajectories with geometry paralleling the population dynamics described in intracortical recordings from motor cortex [1, 2, 29, 30]. As high-gamma activity correlates with neuronal firing rates, these surface recordings effectively sample aggregate spiking across mesoscale cortical territories, making them amenable to similar population-level analyses as traditionally applied to single-unit data [32, 33]. The accessibility of dynamical motor representations from the cortical surface, without penetrating electrodes, extends the dynamical-systems framework to a new recording modality and suggests that the essential geometric structure of motor cortical computation is preserved at the mesoscale while allowing recording from larger cortical territories than are typically accessible with penetrating arrays.

Our density and ablation analyses provide a quantitative framework for understanding why prior surface recordings failed to extract directional information and for guiding the design of future devices. Decoding performance declined with increasing electrode spacing, with the steepest losses occurring above one to two millimeters. This transition provides a direct explanation for why standard clinical arrays, which typically sample at spacings of several millimeters or more, have not supported continuous directional control [23, 26]. Importantly, performance depended specifically on electrode density rather than electrode count, as randomly subsampled channel sets at full density outperformed regularly spaced configurations with equivalent numbers of electrodes. At the same time, spatially informative sites were distributed in unpredictable cortical clusters rather than concentrated in a single focal region of the gyrus, suggesting that broad-area, high-density coverage is essential, as it is likely that smaller arrays at the same pitch would miss critical zones entirely. Redundancy within the current footprint confers robustness to partial channel loss, a property with direct relevance to chronic devices in which individual electrodes may degrade over time.

Beyond the mechanistic findings, these results demonstrate the feasibility of deploying micro-ECoG-based BCIs in a clinically realistic setting. Decoders trained on minutes of data during awake craniotomies supported reliable cursor control within a single operative session, and short-term postoperative implantation yielded highly successful, cross-day performance without retraining, as well as preliminary evidence of cursor control without overt movement. This rapid calibration contrasts with prior ECoG approaches, which have typically required prolonged training or supported only constrained forms of control [6, 19, 21–25]. More broadly, the intraoperative testing paradigm introduced here may help address a persistent bottleneck in brain–computer interface development: the limited number of chronically implanted participants, which constrains the pace at which decoding algorithms can be tested and refined across individuals [34–36]. The ability to train, deploy, and evaluate decoders within a single operative session, across diverse participants and tasks, could accelerate iterative algorithm development and inform the design of future chronic interfaces.

Several important limitations constrain the interpretation of these findings. The cohort was small and heterogeneous, and the intraoperative participants were each limited to a single session with an evolving methodology. The most compelling results, including cross-session generalization and complete task success, came from a single extraoperative case that permitted additional data collection. Furthermore, decoder training relied on joystick-based movements in intact participants, thus the recorded signals likely reflected a combination of motor execution, proprioceptive feedback, and somatosensory reafference, all of which may be altered in people with paralysis. The preliminary finding that one participant maintained cursor control without overt movement provides initial evidence that decoded signals are not entirely execution-dependent, but this result cannot substitute for direct evaluation in the target clinical population. Establishing whether comparable performance can be achieved using attempted or imagined movement in people with paralysis is the central open question for clinical translation. Answering it will also require the development of fully implantable chronic hardware and demonstration of sustained performance over months to years.

Together, these findings redefine what is achievable from the cortical surface. By showing that high-density surface recordings recover the directional population dynamics previously thought to require penetrating electrodes, this work establishes micro-ECoG as both a window onto human motor cortical computation and a scalable platform for brain-computer interfaces. The convergence of a mechanistic account of surface-level population coding, a quantitative framework for array design, and a rapid intraoperative testing paradigm provides a foundation for the next generation of minimally invasive neural interfaces. If evaluation in paralysis, chronic implantation, and long-term stability are demonstrated, high-density micro-ECoG could offer a practical path toward restoring independent communication and computer access for people with severe motor disability.

## 4 Materials and Methods

### 4.1 Participant Cohort, Study Design, and Ethics Approval

The study was approved by the institutional review boards of Johns Hopkins Medicine (IRB00431410) and The Mount Sinai Hospital (IRB Protocol 24-01577), and written informed consent was obtained from all participants. Five participants were enrolled. Four participants undergoing awake craniotomy for tumor resection received temporary intraoperative placement of a 1,024-channel micro-ECoG array over the hand area of motor cortex localized using standard cortical mapping. One participant (P5) underwent an anesthetized craniotomy for tumor resection and received short-term implantation of the 1,024-channel micro-ECoG array over the hand area of motor cortex, localized using standard cortical mapping. For this participant, the micro-ECoG array remained implanted during a clinically determined post-operative monitoring stay of two days.

### 4.2 Experimental Procedure

#### 4.2.1 Intraoperative Recordings

P1-4 underwent awake craniotomies according to routine neurosurgical and anesthesia procedures as detailed in [37]. Participants used the hand contralateral to the side of surgery to perform a 2D target acquisition task with a joystick from a lateral position with the working arm down, moving a screen cursor to sequential designated locations (Fig. 1).

#### 4.2.2 Extraoperative Recordings

A fifth participant (P5) was implanted for a short duration during a two day post-operative monitoring stay in the hospital. This participant completed the same cursor-control task during bedside recordings.

### 4.3 Data Acquisition

Neural activity was recorded using a previously described high-density 1,024-channel micro-ECoG array (Layer 7 Cortical Interface, Precision Neuroscience, New York, NY) [38]. Arrays were made of a flexible polyimide substrate with platinum electrodes spaced at 400 *µ*m pitch, covering approximately 1.5cm^2^ of cortical surface. 1019 central 50 *µ*m and 380 *µ*m diameter contacts contributed to neural features used for decoding, with five larger contacts (500 *µ*m) outside the main grid excluded from analysis (Fig. 1). Signals were acquired using an RHD recording system (Intan Technologies, CA, USA) at 20kHz sampling rate. Task control, decoder inference, and data acquisition were implemented in Python. Neural and behavioral events were synchronized using a shared hardware timebase. In P1-P2, two arrays were placed on the motor strip (Fig. 1). For online decoding in both cases, data only from the array labeled A was used. For P1, it was found offline that array B had superior decoding performance and was therefore used for subsequent analyses. For P2, data from array A was used for all analyses. In P3-P5 a single array was used. Array positions were coregistered to a standard brain anatomy (Fs average brain) based on intraoperative optical imaging and expert anatomical alignment (Fig. 1).

### 4.4 Cursor Control Tasks

Participants performed a joystick-based two-dimensional cursor-control task during sequential recording blocks, each lasting approximately 4 minutes. For participants P1–P3, a center-out paradigm was used. Circular targets appeared sequentially in one of four or eight directions (0°-315° in 45° increments), each placed at a fixed distance from the starting position. For participants P4–P5, targets were presented within a grid to capture continuous movement across a broader range of directions. The grids occupied a fixed volume, with dimensionalities of 4×4 (P4), 12×12, 30×30 (P5 training blocks) and 8×8 (most P5 inference blocks except for a few outliers at 12×12 and 4×4). A trial was considered successful when the cursor entered the target region and remained within it for a predefined dwell period of 0.3 (P4-P5) or 1 second (P1-P3). Trials were terminated if the target was not acquired within either a fixed timeout of 15 seconds (P3) or according to the data collector’s discretion (P4-P5).

Cursor control tasks were completed under one of three different paradigms. During training blocks neural activity was recorded while participants controlled the cursor using a joystick. The velocity of the cursor along the x and y axes was set to be proportional to the instantaneous magnitude of the joystick’s deflection along the respective directions, and the cursor’s pixel position was updated at approximately 60 hz. During typical inference blocks cursor motion was driven in real time by micro-ECoG signals using trained decoders, although participants continued to move the joystick for continuity of experimental conditions. In selected ’imagined’ inference blocks cursor motion was again determined by online decoder output, this time without any joystick present, and participants were instructed to avoid overt hand movement.

For P1 and P2, class-weighted accuracy *A_w_* was calculated as

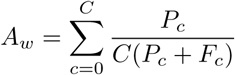

for C total classes, each with *P_c_* correct trials and *F_c_* incorrect trials.

### 4.5 Online Decoding for Closed Loop Cursor Control

Recordings from at least three blocks of approximately four minutes each were used to train a model within a few minutes time, either in the operating room or at bedside. The neural data was downsampled to 1 kHz, filtered to 70 - 150 Hz and further decimated to 100 Hz. For P1-P2, each task block was sequenced into individual trials based on joystick kinematic data that was logged during the experiment. The preprocessed neural data and labeled direction of movement of the cursor in four directions (i.e. up, down, left, and right) during the task trials were used as features for a classification model for P1-P2. The classification model used a convolutional neural network architecture with two fully connected layers applied with activations, pools, and normalizations. During online classification, movement onset was triggered by joystick motion, and a prediction was made using a fixed window of [−400 ms, 200 ms] around movement onset. For P3-P5, the preprocessed neural data and the continuous joystick kinematics were used as features for a convolutional recurrent neural network with a unidirectional gated recurrent units backend. Each of these models was then implemented online for control of the cursor on the task display presented to the patient. In all cases (with one exception, noted below) the patient was instructed to attempt to complete the task using the same physical movements of the joystick, but the cursor was controlled by the decoded neural output. In one session with P5, the patient was instructed to attempt to control the cursor without any hand movements.

In Fig 2 we report Success Rate as the ration of completed tasks to the total number of presented tasks. Completion time measures the duration of completed trials, while throughput captures the overall number of completed tasks per minute, including failed trials.

We did not report regression performance metrics (i.e. *R*^2^, MSE) for the online predictions. Participants instinctively adapted their behavior in response to the model output, forming a closed-loop system that transformed the relationship between joystick tilt and the target cursor motion. Thus direct correlation of cursor and measured joystick motion ceased to be a figure of merit. We instead measured online performance by target completion and acquisition time.

### 4.6 Offline Data Pre-Processing

Channels with impedance *>* 4*M* Ω were excluded. Signals were downsampled to 1kHz, re-referenced to a common average, and notch-filtered at 60 Hz and harmonics to remove line noise. HGA features were extracted by band-pass filtering (70-170 Hz (P1-P2) or 70–150 Hz (P3-P5), 5th-order Butterworth), computing the analytic amplitude, and applying an exponential moving average (EMA; *α* = 0.3). Features were normalized online using a running zero-mean transform and decimated by a factor of 10 to match the temporal resolution of joystick kinematics. For P5, the band-pass range was reduced to 70–115 Hz to further suppress line-noise contamination.

When relevant, the recordings were segmented into trials using a speed-based thresholding procedure. The cursor speed was computed as 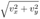 and smoothed with a centered moving average of 50ms. Trials were defined as continuous epochs where smoothed speed exceeded an absolute threshold (0.15 a.u.) including 400ms before and after the epoch to include the peri-movement context. Overlapping epochs were not merged. Trials were padded to enable fixed-length batching. Direction was assigned per trial by computing the mean velocity over the “active” portion of movement as defined as the top 40% of samples. The vector’s angle was converted to degrees per the convention of the study (0°, 45°, 90°, 135°, 180°, 225°, 270°, or 315°) and binned to the nearest of those targets. It is worth noting that while both the center-out and grid task were binned to the same set of angles, the fidelity of this abstraction is more limited for the grid task.

### 4.7 Pseudo-online Regression Analysis

The processed HGA data (now sampled at 100Hz) was organized into overlapping temporal windows (stride 200ms), to emulate the batch-wise real time processing used in closed-loop experiments. Each HGA window was linked with the 2D joystick position value corresponding to the last sample per window to generate input–output training pairs. This resulted in an input dataset dimensionality of ℝ*^X·Y·F·T·N^* and an output dataset dimensionality of ℝ^2·*N*^, where *X* and *Y* define the respective dimensionality of the ECoG grid in electrodes, *F* quantifies the number of features per electrode (1 for all networks using HGA features), T defines the number of prior time samples included in each window (350-700 depending on participant), and N is the number of windows extracted from each recorded block. Each window is passed singularly through a CRNN implemented in PyTorch. For the training, window order was shuffled, but for testing the points were evaluated sequentially.

The model first applied a 3-D convolution to collapse an input layer in ℝ*^X·Y·F·T^* to a lower dimensional latent space. This was followed by a Bi-LSTM to capture temporal dynamics. Then the final output of the LSTM was projected through a fully connected layer to predict instantaneous velocity (*V_x_, V_y_*). Because each velocity value is calculated using a single window of HGA data, this analysis remains causal because the bi-directional LSTM layer of the LSTM only operates on T HGA samples, and a new velocity value is generated for every 200ms of data collected. Dropout was applied at both stages to reduce overfitting. The network was trained using Adam optimization on a mean squared error (MSE) loss function.

A model was trained for each participant using the first three blocks of labeled data, while the remaining labeled data block was used for hyperparameter optimization and evaluation. Predictions were compared against ground-truth joystick velocities for both x- and y-components. Final analyses demonstrated the ability of high-gamma features recorded from the 1,024-channel µECoG array to accurately reconstruct cursor velocity trajectories.

To assess the quality of the regression models we calculated the coefficient of determination (*R*^2^) between the recorded cursor velocity and our pseudo-online results.

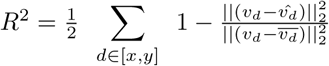

For recorded velocity *v*[*t*] of length T with mean *v̄*, range Δ*V* and predicted velocity *v̂*[*t*].

We also report percent mean square error (%MSE) as

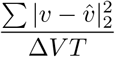

### 4.8 Hyperparameter search

We performed a two stage participant-specific hyperparameter optimization using the Optuna framework [39]. Stage 1 explored 100 configurations using Tree-structured Parzen Estimator sampling with median pruning for early stopping. The search space included the time captured per input sample (0.5s-4s), CNN architecture parameters (kernel sizes: 4–10 spatial, 20–60 temporal; kernels: 2–8), LSTM parameters (hidden units: 128–800; layers: 2–6), dropout rates (0.2–0.7 CNN, 0.1–0.5 RNN), and optimization parameters (learning rate: 10^−5^–10^−3^; weight decay: 10^−5^–10^−2^). Stage 2 validated the top 10 configurations with 5 runs each. Optimization was based on the median *R*^2^ scores from the 5 runs to ensure robust performance across random initializations. All models were trained for 100 epochs using the Adam optimizer.

### 4.9 Electrode Density Analysis

A CRNN model was trained for each participant and electrode subset configuration. Per-participant optimized hyperparameters were used, with the exception of the spatial parameters, which were defined to restrict the model to temporal convolution. To control for the intrinsic stochasticity of network training, each configuration was trained and evaluated three times. Nonlinear least squares was used to fit an exponential decay model *F*(*d*) with constants *C* to measured *R*^2^ and electrode pitch *d* data for each participant.

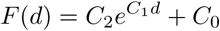

We calculated 95% confidence intervals by assuming a t-distribution for the residuals of each polynomial fit.

### 4.10 Ablation analysis

To characterize the spatial distribution of decoding-relevant neural activity across the micro-ECoG array, we performed a multi-scale spatial ablation analysis. Using the same convolutional-BiLSTM architecture described above with the spatial convolutional kernel reduced to 1×1 to prevent spatial mixing across electrodes, we systematically zeroed contiguous *N x N* electrode tiles (*N* = 2, 3, 4, 5, 6) from the 32×32 grid during held-out validation and quantified the resulting change in decoding performance. Tile importance was defined as the fractional drop in *R*^2^ relative to the unablated baseline: Δ*R*^2^*_norm_* = (*R*^2^*_baseline_* − *R*^2^*_masked_*)*/R*^2^*_baseline_*, computed separately for each velocity component and summed Δ*R*^2^*_x_*_+*y,norm*_ = Δ*R*^2^*_x,norm_* + Δ*R*^2^*_y,norm_*. For per-electrode importance, tile-level scores were aggregated to individual channels by averaging the Δ*R*^2^ values across all overlapping tiles containing each electrode. Using the (x, y) coordinates from the probe map, per-channel importance was rendered as spatial heatmaps and used to generate ranked lists of informative channels.

To test whether electrode importance was spatially clustered beyond what would be expected by chance, we computed Moran’s I — a measure of spatial autocorrelation — on each per-electrode importance map using queen-contiguity (8-neighbor) weights on the 32×32 grid. Significance was assessed via permutation testing: the observed importance values were randomly shuffled across occupied grid positions 9,999 times, preserving the marginal distribution while destroying spatial structure. A one-sided p-value was computed as the fraction of permuted Moran’s I values equal to or exceeding the observed statistic.

### 4.11 µECoG Directionality Analyses

For subsequent population summaries, HGA was normalized (z-score relative to 0.0-0.3 s prior to defined movement onset), then averaged within each direction to yield trial-averaged mean and SEM time courses per channel. *ChannelXTime* heatmaps and time courses were plotted to visualize directional tuning and temporal dynamics.

#### 4.11.1 PCA to UMAP Dimensionality reduction

To visualize the low-dimensional geometry of population neural activity during movement, we computed dynamic latent-space trajectories using a two-stage dimensionality reduction pipeline. Neural data were first partitioned into overlapping sliding windows (width = 150 ms, step = 40 ms) spanning the analysis epoch (0–1.3 s post-movement onset). Within each window, the HGA of every channel was averaged across time, yielding a feature matrix of shape (*N* × *W* × *C*), where *W* is the number of windows. All (*N* × *W*) window-trial observations were then projected into a principal component space using 100-250 PCs, retaining the leading components to explain the majority of the variance. The resulting scores were embedded into three dimensions using Uniform Manifold Approximation and Projection (UMAP; *n* neighbors = 50, min dist = 0.01, cosine metric). The embedding was reshaped back to (*N*, *W*, 3), producing a per-trial trajectory through the three-dimensional UMAP space. Direction-averaged trajectories were smoothed with a Gaussian filter (*σ* = 1 windows) and plotted as trajectories to convey the mean path and inter-trial variability for each reach direction. This visualization was used for qualitative assessment only; all quantitative decoding and separability metrics were computed in PCA space (see below).

To quantify the time course of directional information in the neural population, we performed time-resolved classification using a sliding-window approach. At each window center, the channel-wise mean HGA vector (one scalar per channel) served as the feature representation for each trial. Classification was performed using regularized linear discriminant analysis (LDA). A five-fold stratified cross-validation procedure was used, with principal component analysis (PCA) fitted on the training fold only at each window to prevent information leakage from test trials. Classification accuracy was averaged across folds to produce a single accuracy value per time window.

To dissociate the relative contributions of spatial (cross-channel) patterns and temporal dynamics to directional encoding, we applied two complementary permutation controls to the neural data before repeating the time-resolved decoding analysis. For each trial independently, the time axis was randomly permuted using a single permutation shared across all channels and the channel axis was randomly permuted using a single permutation shared across all time points. To establish a statistical significance threshold, we repeated the full decoding pipeline with randomly permuted trial labels (*n* = 200 permutations). At each time window, the 2.5th and 97.5th percentiles of the resulting null accuracy distribution defined a 95% confidence interval for chance-level performance. The theoretical chance level for eight-class direction classification is 12.5% (1/8).

As a complementary, classifier-free metric of directional separability, we computed a time-resolved between/within distance ratio in PCA space. We quantified geometric separability of direction-specific trajectories using a ratio of inter-condition distance to within-condition dispersion, conceptually analogous to classical between/within-class separation metrics used in discriminant analysis and clustering, and consistent with prior population-dynamics analyses of motor cortical trajectories [1, 2]

The separation index was defined as:

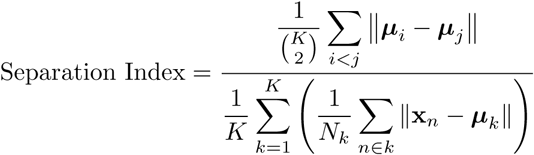

or

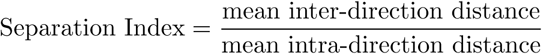

This metric was computed for the original data and both permutation controls using the same sliding-window and PCA parameters as the decoding analysis.

## Acknowledgments

We gratefully acknowledge clinical coordination from Emily Svendsen, Kevin Buttet, and Kimberly Agosto from Mount Sinai Health System. We also acknowledge clinical coordination and figure illustration from Meghan Hildreth and Jaime Chong from Precision Neuroscience.

## Funding

This work was supported in part by philanthropic funding from Dr. Charles Fields. The donor had no role in study design, data collection, data analysis, data interpretation, or writing of the manuscript.

## Competing interests

R.A., Y.W.B, J.S.M., M.M., J.D., K.B., A.S., E.H., E.G., J.B., F.C., M.L., C.H.M., J.S., B.I.R., are current or former employees or advisors of Precision Neuroscience and own stock in the company.

